# Yeasts dominate soil fungal communities in three lowland Neotropical rainforests

**DOI:** 10.1101/138057

**Authors:** Micah Dunthorn, Håvard Kauserud, David Bass, Jordan Mayor, Frédéric Mahé

## Abstract

Forest soils typically harbour a vast diversity of fungi, but are usually dominated by filamentous (hyphae-forming) taxa. Compared to temperate and boreal forests, though, we have limited knowledge about the fungal diversity in tropical rainforest soils. Here we show, by environmental metabarcoding of soil samples collected in three Neotropical rainforests, that Yeasts dominate the fungal communities in terms of the number of sequencing reads and OTUs. These unicellular forms are commonly found in aquatic environments, and their hyperdiversity may be the result of frequent inundation combined with numerous aquatic microenvironments in these rainforests. Other fungi that are frequent in aquatic environments, such as the abundant Chytridiomycotina, were also detected. While there was low similarity in OTU composition within and between the three rainforests, the fungal communities in Central America were more similar to each other than the communities in South America, reflecting a general biogeographic pattern also seen in animals, plants, and protists.

## 1 BACKGROUND

Fungi are microbial eukaryotes that are constituents of most ecosystems, where they fulfil diverse ecosystem functions such as saprotrophy, mutualism, and pathogenesis/parasitism (Blackwell, 2011; Peay *et al.*, 2016; Treseder and Lennon, 2015). They have different growth forms, ranging from unicellular organisms (James *et al.*, 2006; Jones *et al.*, 2011) to widespread mycelial networks that can occupy several square kilometres (Smith *et al.*, 1992). Filamentous growth forms typically dominate terrestrial ecosystems (Peay *et al.*, 2016; Treseder and Lennon, 2015). However, throughout the fungal tree of life, especially in the Ascomycota and Basidiomycota, many groups have evolved unicellular yeast morphologies either throughout their lifecycle or only under certain aqueous conditions (Botha, 2011). Some, like the Saccharomycetes (hereafter referred to as Yeasts) have only unicellular forms, while other taxa can switch between filamentous and unicellular morphologies depending on environmental conditions. Yeasts are known to dominate fungal communities in liquid or stressful environments such as the rumen of farms animals and deer (Kittelmann *et al.*, 2013), floral nectaries such as the bumble-bee-pollinated *Helleborus foetidus* (Herrera *et al.*, 2010), and deep-ocean water and sediments (Bass *et al.*, 2007).

While temperate, boreal and arctic soil fungi communities have been examined in detail using environmental high-throughput sequencing methodologies (e.g., Clemmensen *et al.*, 2013; Mundra *et al.*, 2016; O’Brien *et al.*, 2005; Tedersoo *et al.*, 2014), the diversity of soil fungal communities in the tropics is relatively rarely sampled. For example, Tedersoo *et al.* (2014) found the Agaricomycotina to be dominating the fungal communities in their Neotropical forests sites and that Yeasts comprised less than 1% of their operational taxonomic units (OTUs). Likewise, Kivlin and Hawkes (2016) sampling of monoculture plots and secondary forests in La Selva Biological Station, Costa Rica, and Peay *et al.* (2013) sampling of forests in Amazonian Peru found the Agaricomycotina to be dominating. Peay *et al.* did note they found many Yeasts, which accounted for almost 25% of their sequencing reads.

To more thoroughly analyse Neotropical rainforest soil fungal communities, we extensively sampled three Central and South American forests and deeply sequenced using Illumina MiSeq. The hyperdiverse protist communities were recently analysed from these same samples, and a large proportion of OTUs were found to be highly dissimilar to those in reference sequence databases, to be dominated by parasites, and to exhibit extremely high heterogeneity between spatially disjunct samples even within the same forests (Mahé *et al.*, 2017). Using the same DNA extracted from the soils, we asked what is the dominant fungal group(s) and if the fungal communities exhibit comparable patterns of biodiversity.

## 2 MATERIALS AND METHODS

### (a) Sampling, sequencing, and clustering

All codes used here can be found in HTML format (supplementary file 1). Details about sampling and permits can be found in Mahé *et al.* (2017). Briefly, 279 soils samples were taken in a variety of lowland Neotropical forest types in: La Selva Biological Station, Costa Rica; Barro Colorado Island, Panama; and Tiputini Biodiversity Station, Ecuador. DNA was extracted, and every two samples were combined. The samples were first amplified for the hyper-variable V4 region of 18S rRNA locus following Mahé *et al.* (2015a) with general eukaryotic V4 primers (Stoeck *et al.*, 2010), then amplified with sample-specific tags and Illumina MiSeq’s sequencing adapters. Illumina MiSeq sequencing used v3 chemistry.

Fastq files were assembled with PEAR v0.9.8 (Zhang *et al.*, 2014) using default parameters and converted to fasta format. Paired-end reads with both primers and no ambiguous nucleotides were retained with Cutadapt v1.9 (Martin, 2011). Reads were dereplicated into strictly-identical amplicons (that is, reads were merged at 100% similarity and to which an abundance value can be attached) with VSEARCH v1.6.0 (Rognes *et al.*, 2016), and clustered into OTUs with Swarm v2.1.5 (Mahé *et al.*, 2015b) using d = 1 with the fastidious option on. Chimeric OTUs were identified and removed with VSEARCH *(de novo* search). Low abundant OTUs were discarded only if they included <2 reads, and were sequenced from only one sample, and were <99% similar to accessions in the Protist Ribosomal Reference (PR2) database v203 (Guillou *et al.*, 2013). Following Adl *et al.* (2012), the PR2 database contains 21,083 fungal references.

### (b) Analyses

Stampa plots (Mahé, 2016) were made to show the distribution of the number of reads and OTUs per similarity value to their best match in the PR2 database. Taxonomic assignment of the amplicons and OTUs used VSEARCH’s global pairwise alignments with the PR2 database. Assignment used the best hit, or co-best hits, in the reference database as reported by VSEARCH. The R package Vegan (Oksanen *et al.*, 2013) was used to analyse frequency count data derived from OTU clustering. Different functions of Vegan were called to randomly subsample our samples (rrarefy function) and to estimate and compare species compositions, using Bray-Curtis distance and NMDS ordination (monoMDS function). Figures were made using the R Statistical Environment (R Core Team, 2016) and ggplot2 (Wickham, 2009).

## 3 RESULTS AND DISCUSSION

### (a) Read and OTU characteristics

From the soil samples collected in the three Neotropical rainforests, a total of 44,430,656 cleaned reads comprising 17,849 OTUs were assigned to the fungi (for the OTU table see supplementary file 2). With our use of the Swarm clustering method, 99.9% of the OTUs had radii with >97% similarity (figure 1), which is a widely used global clustering threshold in other 18S rRNA fungal environmental sequencing studies. About 91.2% of the total reads and OTUs had a maximum similarity of ⩾95% to references in the PR2 database, and almost 97.6% of them were ⩾80% similar to the references (figure 2). This pattern of mostly high similarities to PR2 references also occurs in most of the different fungal subtaxa (supplementary figures 1 and 2). In contrast to these high fungal similarities from the same soil samples, only 14.7% of the protists had a maximum similarity of >80% to PR2 references (Mahé *et al.*, 2017). Neo-tropical rainforest fungi are therefore better characterised than other microbial eukaryotes or at least have more closely related relatives that have already been sequenced.

**Fig. 1.**
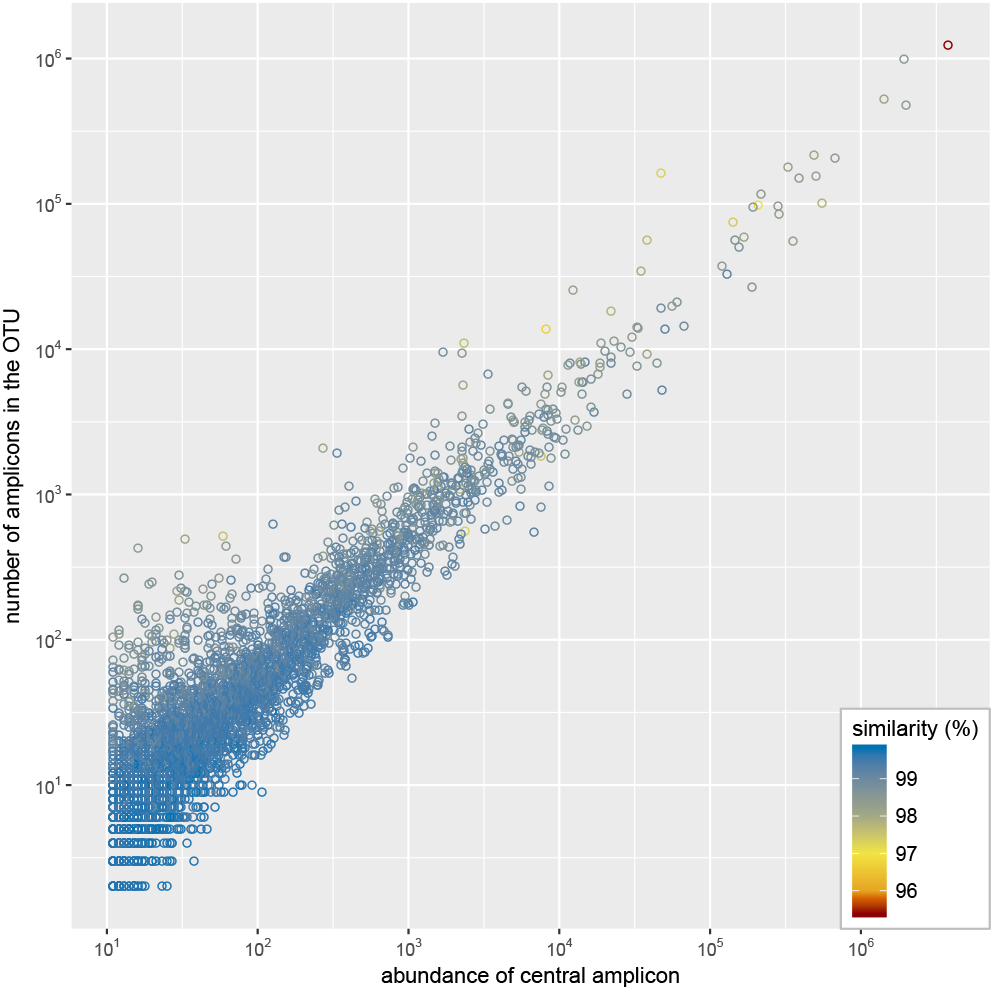
The radius (minimum % similarity between the OTU centroid and any other member of the OTU) of Swarm OTUs of the fungal V4 data. The number of amplicons in each OTU was plotted against the abundance value of the OTUs’ centroids (most abundant amplicon). More than 99.9% of the OTUs have radii >97% similarity.

**Fig. 2.**
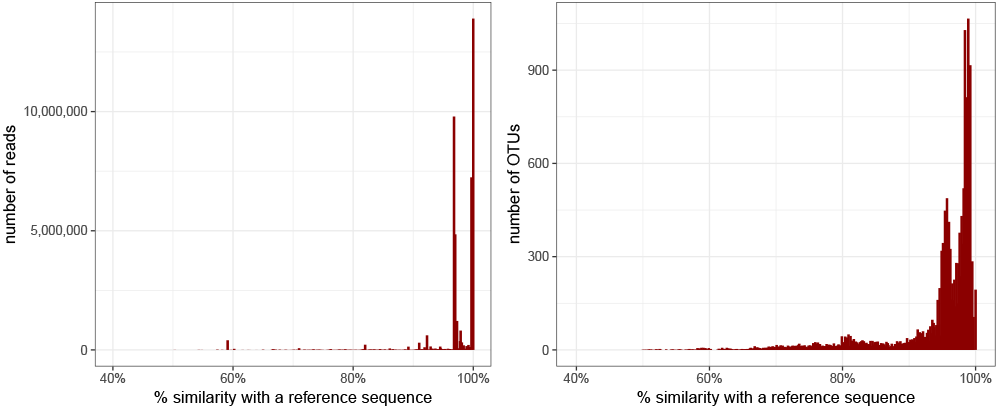
Similarity of fungi reads and OTUs to the taxonomic reference database. Most of the reads and OTUs were ⩾80% similar to references in the PR2 database. See supplementary figures 1 and 2 for these similarities within different fungal subclades.

### (b) Yeasts dominate Neotropical soil fungal communities

A dominating part of the fungal diversity in the three Neotropical rainforests were the Yeasts and other unicellular-forming fungi (figure 3, supplementary file 2). The Saccharomycotina (in the Ascomycota) were the most abundant fungi for both reads (63.8%) and OTUs (37.0%). This proportional higher abundance of Yeast reads may indicate that they are represented by a high biomass or number of cells in the soil, although in a recent study the proportion of fungal reads to OTUs was largely congruent (Khomich *et al.*, 2017). Agaricomycotina (in the Basidiomycota) was also highly represented (21.0% of the reads and 20.4% OTUs), but this group was also mainly composed of OTUs with taxonomic affiliation to unicellular or dimorphic forming tremellomycetes (Liu *et al.*, 2015), such as Trichosporon spp. Tremellomycetes accounted for 79.7% of the reads, and 94.0% of the OTUs of the Agaricomycotina. Some other fungal groups, including Pucciniomycotina and Ustilaginomycotina were also largely represented by unicellular anamorphic fungi, suggested by a majority of matches towards Sporobolomyces.

**Fig. 3.**
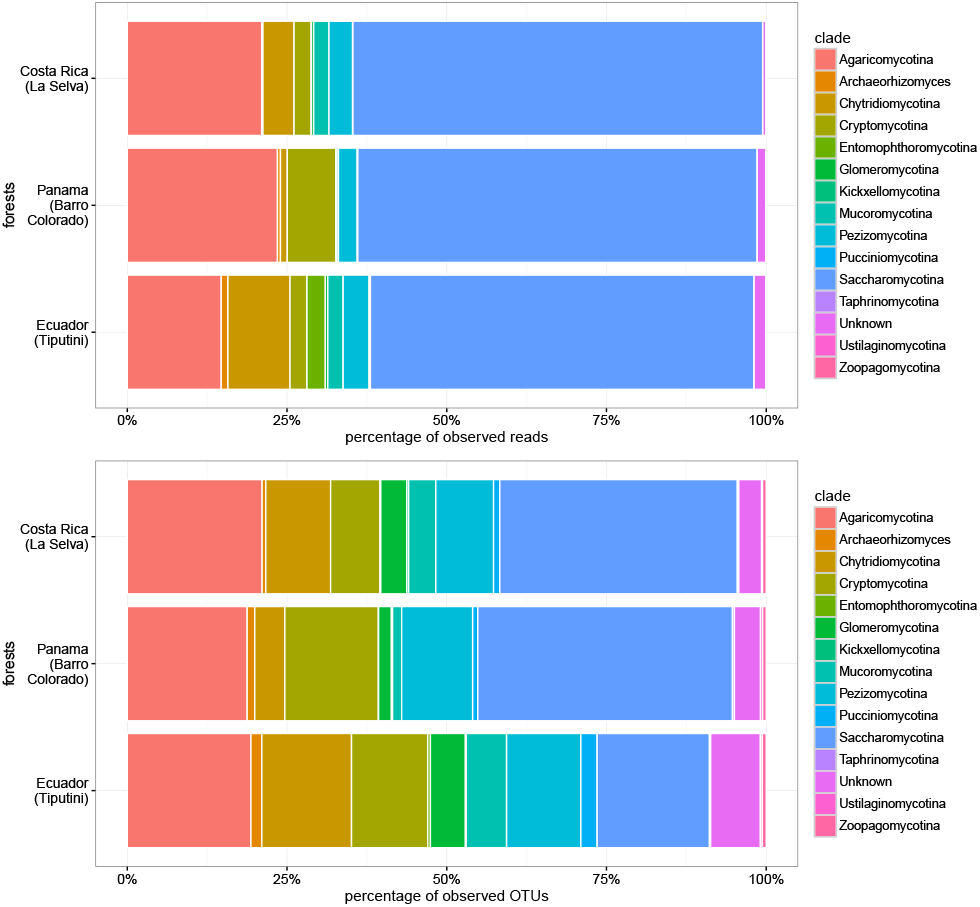
Taxonomic identity and relative abundances of fungi reads and OTUs. Each taxa shown represent at least >0.1% of the total data. Taxonomic assignment is to the fifth level in PR2 reference database, except for Archaeorhizomyces.

Proportionally fewer Yeasts were detected in previous environmental sequencing studies of fungal communities in Neotropical forests (Tedersoo *et al.*, 2014; Kivlin and Hawkes, 2016; Peay *et al.*, 2013). Our results may differ from these studies for a number of reasons. First, we employed very deep Illumina MiSeq sequencing on a large number of independent samples. Second, we amplified the V4 region rather than the ITS region like Tedersoo *et al.* (2014) and Peay *et al.* (2013), or LSU region like Kivlin and Hawkes (2016). Although ITS primers are widely used for fungi (Blaalid *et al.*, 2013), 18S rRNA primers have also been used to effectively characterize fungal communities (Bass *et al.*, 2007; Richards *et al.*, 2015), but all primers have biases. Third, we used local clustering thresholds to produce the fine-grained Swarm OTUs rather than global clustering thresholds. Many of these fine-grained OTUs would have been otherwise lumped together with clustering methods that employed a global clustering threshold of 97% similarity. And fourth, we sampled primary forests regardless of tree species, rather than sampling forests with ectomycorrhizal plant species like Tedersoo *et al.* (2014) and secondary and cultured forests like Kivlin and Hawkes (2016).

Yeasts could be dominating the fungal communities in the three Neotropical rainforests because of the predominantly aqueous conditions in the soils due to frequent (sometimes daily) flooding leading to inundated or near-saturated soil conditions (Silver *et al.*, 1999). Seasonal and prolonged anoxia is known to structure microbial communities (Pett-Ridge and Firestone, 2005; Pett-Ridge *et al.*, 2006) and to alter ecosystem functions including nitrogen and carbon cycling (Davidson *et al.*, 2000; Dubinsky *et al.*, 2010; Turner *et al.*, 2015; Waldrop *et al.*, 2000). The variable oxygen concentrations could also create steep gradients in redox within centimetre or millimetre scales that permit coexistence of numerous aerobic and anaerobic microorganisms within close proximity (Sexstone *et al.*, 1985). The frequent anoxia therefore would be highly influential over the Neotropical soils examined here, and such unicellular fungal morphologies may be advantageous in such a saturated environment, particularly when simple sugars are available due to high turnover of soil carbon. In addition to soil-inhabiting Yeasts and unicellular-forming fungi, there are several other habitats in the tropical rainforests that could harbour these taxa that then wash down onto surface soils (Botha, 2011). For example, Yeasts and unicellular-forming fungi are known to inhabit animal rumens, lichen thalli, and plant leaves, fruits, flowers, and wood, and to use insect vectors to move between them (Botha, 2011; Abranches *et al.*, 1998; Li *et al.*, 1985; Rosa *et al.*, 2007; Spribille *et al.*, 2016). In the vertical arrangement of forests, from soils to tree canopies, numerous flowers and fruits will exist in different stages of ripening, representing numerous ecological niches for the growth and promotion of unicellular fungi (Morais *et al.*, 2006).

The proportional abundance of these detected Yeasts raise a question: What are they all doing in the tropical soils? They are presumably breaking down simple sugars near plant roots and fallen fruits, or degraded lignin elsewhere in the soils (Botha, 2011; Rodrigues *et al.*, 2006). During this breakdown of carbon compounds, the most common pathway would be anaerobic fermentation (Rodrigues *et al.*, 2006), especially given the anoxic conditions caused by water inundation. Even in the absence of anoxic conditions, anaerobic-and aerobic-fermentation could take place via the well characterised Crabtree effect that produces ethanol (De Deken, 1966). Better understanding of the biogeochemical role of unicellular fungi in tropical soils would help parameterise how carbon and nutrient cycles and soil respiration respond to saturating conditions (Angert *et al.*, 2015). For instance, slowed transport processes and the associated accumulation of decomposition products in water saturated soils present distinct enzymatic challenges to the degradation of soil carbon by phenol oxidases (Limpens *et al.*, 2008)—enzymes produced by a wide range of fungi (Burke and Cairney, 2002; Hammel, 1997) and other microbial organisms. The consequences of these processes include a switch from high soil CO_2_ efflux to the atmosphere under oxic conditions to the transport of dissolved carbon in the hydrological system under anoxic conditions (Angert *et al.*, 2015).

Numerous other fungi were also found in the Neotropical rainforest soil samples. The Chytridiomycotina, a large saprotrophic and parasitic radiation also found in aqueous conditions, were likewise prevalent especially in terms of OTU richness (1,671 OTUs). Many chytrids are likely not amplified by ITS-based studies due to primer mismatches. Although most of the detected fungi are typically found in aquatic environments, filamentous terrestrial fungi were also detected, including members of Agaricomycotina, Archaeorhizomyces, Glomeromycotina, Mucoromycotina, and Pezizomycotina. The arbuscular mycorrhizal Glomeromycotina, globally distributed in terrestrial habitats (Davison *et al.*, 2015), had a high OTU richness (674 OTUs) but made up a relatively small proportion of the reads. The spores and coenocytic hyphae of Glomeromycotina, which may be cryptically sexual (Halary *et al.*, 2011; Corradi and Brachmann, 2017) and may include numerous genetically different nuclei, complicate and may exaggerate richness estimates. The most abundant OTUs in the Ascomycete Pezizomycotina were affiliated with perithecial Sordariomycetes, which are widespread decomposers of plant litter and dung, but some are common parasites.

Within the Mucoromycotina, we obtained matches to various taxa with different ecologies, including Mortierella and Umbelopsis, assumed to be saprotrophs, and Endogone, a plant root symbiont. We detected 128 OTUs affiliated to the newly described Archaeorhizomycetes (Rosling *et al.*, 2011) that are nested within the Taphrinomycotina (Ascomycota). This fungal group is associated with plant roots and has mainly been detected in forests in the Northern Hemisphere; but our data, together with Tedersoo *et al.* (2014), show that it also is widespread in tropical forests. Additionally, 108 OTUs were assigned to the Zoopagomycotina, which are parasites of other fungi or soil invertebrates, and 32 OTUs were assigned to the Atractiellomycetes (Pucciniomycotina), which form mycorrhizal associations with Neotropical orchids (Kottke *et al.*, 2010).

### (c) Neotropical soil fungi follow general eukaryotic biogeographic patterns

Fungal community diversity as estimated by the Jaccard similarity index showed that there was high heterogeneity in the OTU composition between samples. This high heterogeneity (that is, low number of shared OTUs) occurred both within and between forests (figure 4, supplementary figure 3). Although the Jaccard similarity values were low on average among all samples (0.0111), they were higher, in our recalculation of Mahé *et al.’s* data, than the average for protists from the same samples (0.0069). Even with this low similarity, non-metric multidimensional scaling (figure 5, supplementary figure 4) and Bray-Curtis dendrograms (supplementary figure 5) showed fungal communities in Costa Rica and Panama to be slightly more similar to each other than the fungi in Ecuador. This difference between Central American and South American fungi was also found in protists (Mahé *et al.*, 2017) and in animals and plants (Gentry, 1991); that is, we found a general pattern from the microbial to the macro-organismic levels in the eukaryotes.

**Fig. 4.**
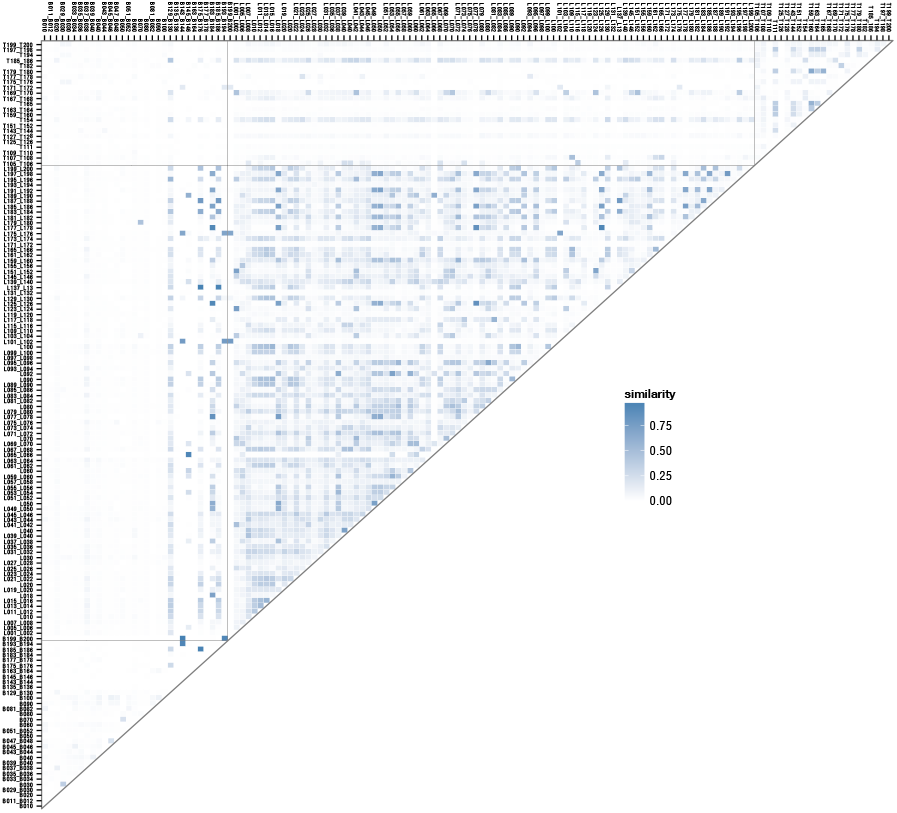
Jaccard similarity index of fungi OTU composition differences between soil samples. See supplementary figure 3 for sample names.

**Fig. 5.**
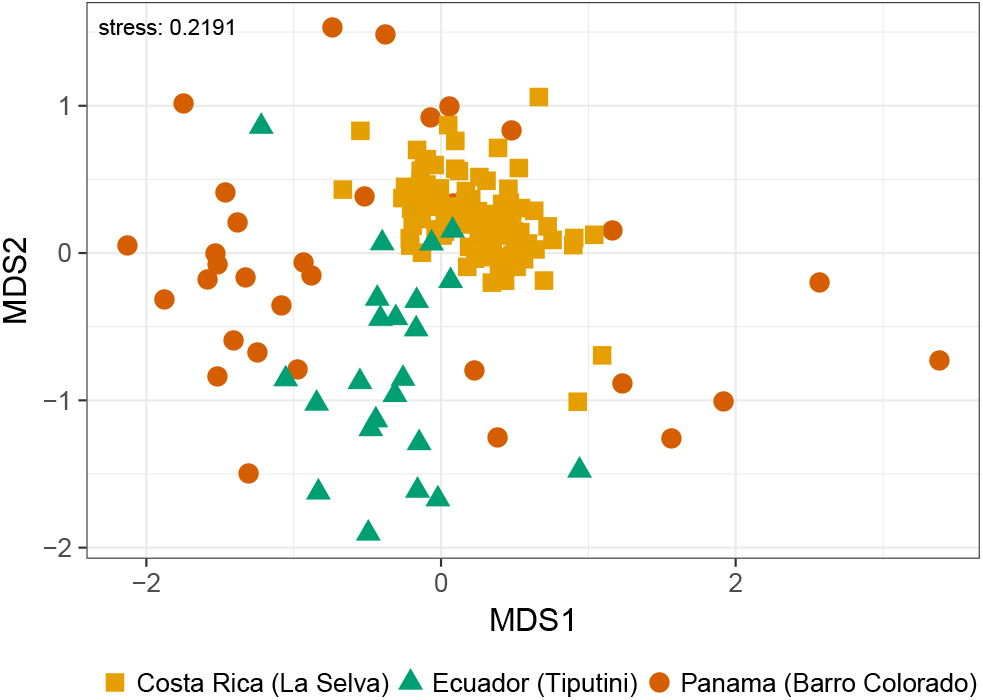
Non-metric multidimensional scaling of fungi OTU composition differences between soil samples. See supplementary figure 4 for sample names.

## 4 CONCLUSION

Similar to fungal communities in some aqueous environments (Kittelmann *et al.*, 2013; Herrera *et al.*, 2010; Bass *et al.*, 2007), Yeasts dominated the communities in three Neotropical rainforests. Frequent anoxic soil conditions caused by heavy rains in these forests could create ideal conditions for unicellular fungi. Dominance by Yeasts suggests tropical forests are not only persistent carbon sinks owing to rapid tree growth (Pan *et al.*, 2011), but also enormous fermentation tanks anaerobically processing carbon because of the fungi.

## Data accessibility

The data analysed in this study are available at GenBank’s Sequence Read Archive under BioProject number SUB582348.

## Authors’ contributions

M.D. and F.M. conceived the project. M.D., J.M., and F.M. collected the data. M.D., H.K., D.B., J.M., and F.M. performed the analyses and wrote the paper.

## Acknowledgements

We thank T. Siemensmeyer for help in sample collecting.

